# A CROP-seq screen of histone modifying enzymes reveals histone demethylase Kdm5c regulates inflammatory macrophage activation

**DOI:** 10.1101/2025.10.30.685537

**Authors:** Femke Horstman, Guillermo R. Griffith, Francesco Vinciguerra, Ricky Siebeler, Cindy van Roomen, Tatyana Kuznetsova, Judith C. Sluimer, Gerard Pasterkamp, Michal Mokry, Menno P.J. de Winther, Koen H.M. Prange

## Abstract

**Background:** Macrophages adopt activation states along a spectrum from pro- to anti-inflammatory, enabling appropriate responses to pathogens and environmental cues. Dysregulated inflammatory macrophage activation contributes to diseases including sepsis, rheumatoid arthritis, cancer, and atherosclerosis. Epigenetic processes such as DNA methylation and histone modification prime macrophages for activation, and several histone modifying enzymes (HMEs) have been implicated in this regulation.

**Objective:** To systematically identify histone modifying enzymes that regulate inflammatory macrophage activation.

**Methods:** We performed a CRISPR knockout screen with single-cell RNA-seq readout (CROP-seq) targeting 92 macrophage-expressed HMEs in immortalized LPS-activated mouse bone marrow-derived macrophages (BMDMs). The resulting single-cell transcriptomes were analyzed to identify significant perturbations. *Kdm5c* was selected for experimental validation in mouse BMDMs, and its expression pattern was compared with macrophage subsets from human atherosclerotic plaques using scRNA-seq data.

**Results:** The CROP-seq screen identified *Prmt6, Carm1, Kat2b*, and *Kdm5c* as top regulators of inflammatory macrophage activation. Validation in a KO cell line revealed loss of *Kdm5c* suppressed inflammatory tone at baseline but led to an exaggerated transcriptional response to LPS stimulation, indicating a role for Kdm5c in balancing tonic and inducible activation. A weighted gene module derived from *Kdm5c*-deficient macrophages was enriched in inflammatory macrophages in human atherosclerotic plaques.

**Conclusion:** Our findings demonstrate the value of CROP-seq screening to dissect the epigenetic control of macrophage activation. We also identify Kdm5c-mediated histone demethylation as a key mechanism modulating inflammatory macrophage activation.

## Introduction

Macrophages respond to pathogens and environmental cues by adopting activation states along a spectrum from pro- to anti-inflammatory(1). Precise control of these states is critical, as aberrantly activated macrophages drive both acute conditions such as bacterial sepsis and chronic diseases including rheumatoid arthritis, cancer, and atherosclerosis(2-7)

Macrophage activation is characterized by the stepwise opening of chromatin(8), mediated by signal-dependent transcription factors that initiate gene programs specific to distinct activation states (e.g. inflammatory or fibrotic). These responses are epigenetically primed through processes such as DNA methylation, histone modification, and promoter-proximal RNA polymerase II stalling(9). Several histone modifying enzymes (HMEs), including HDAC3(10) and KDM6B(11-13), have been shown to modulate macrophage phenotype.

In atherosclerosis—the leading cause of cardiovascular disease(14)—monocytes infiltrate the vessel wall to clear accumulating lipids. Upon lipid uptake, they differentiate into macrophages and subsequently into foam cells(15, 16). These lipid-laden cells can undergo necrosis, releasing their contents and perpetuating inflammation, which attracts further immune cells. This feed-forward process promotes plaque growth and destabilization, ultimately predisposing to rupture and thrombosis(2, 3).

Inflammatory macrophages are key determinants of plaque (in)stability(17). The clinical relevance of targeting inflammation in atherosclerosis, on top of optimal lipid-lowering therapy, was demonstrated by trials such as CANTOS(18). We therefore set out to identify HMEs that modulate inflammatory macrophage activation.

To systematically profile these epigenetic regulators, we performed a CRISPR knockout screen with single-cell RNA-seq readout (CROP-seq)(19, 20) targeting 92 macrophage-expressed HMEs in immortalized mouse bone marrow–derived macrophages (BMDMs). This screen identified *Prmt6, Carm1, Kat2b*, and *Kdm5c* as top hits influencing inflammatory activation. We then validated the effect of *Kdm5c* knockout in mouse BMDMs and found that while *Kdm5c* loss suppressed baseline inflammatory tone, it enhanced the transcriptional response to LPS stimulation, indicating a dual role in balancing tonic and inducible activation.

Finally, to place these findings in a disease context, we compared the *Kdm5c*-dependent gene program and transcription factor networks with macrophage subsets from human atherosclerotic plaques. The *Kdm5c*-deficient signature and its NF-κB–STAT3 regulatory network were enriched in inflammatory foamy macrophages, implicating KDM5C as an epigenetic modulator of inflammatory macrophage states relevant to plaque inflammation and instability.

Together, these findings demonstrate the utility of an HME-focused CROP-seq approach to dissect the epigenetic regulation of macrophage activation and identify KDM5C-mediated histone demethylation as a key mechanism that modulates inflammatory macrophage activation.

## Methods

### iBMDM cells

Immortalized mouse bone marrow derived macrophages (iBMDM)s were generated as described(21) from a Rosa26-Cas9-IRES-EGFP male mouse (Jackson Lab stock No: 028555). iBMDM were cultured in DMEM high glucose, supplemented with 10% FBS, 1% Pen/Strep, 2 mM L-Glutamine, and 17 ng/ml M-CSF.

### ER-Hoxb8 cells

ER-Hoxb8 cells were generated as described (22) from Rosa26-Cas9-IRES-EGFP positive mouse myeloid progenitor cells. ER-Hoxb8 transduced progenitors were cultured in DMEM, supplemented with 10% FBS, 1% Pen/Strep, 2mM L-Glutamine, 20 ng/ml GM-CSF, and 0.5 µM ß-estradiol, and were differentiated to macrophages as needed in DMEM supplemented with 10% FBS, 1% Pen/Strep, 20 ng/ml L-Glutamine, and 17 ng/ml M-CSF for 7 days.

### CROP-seq

CROP-seq was performed as described(19) with the inclusion of a guide enrichment PCR as described(20). Guide sequences were sourced from the GECKOv2 database(23) and ordered as custom oligos. CROP-seq backbone vector was ordered form Addgene (Plasmid #86708). Briefly, we transduced immortalized mouse BMDM with the pooled HME CROP library and let the cells grow in the presence of 4 mg/ml puromycin for 10 days to select for successfully transduced cells and allow time for gene-editing. After, we added 10 ng/ml LPS for 3 hours to stimulate inflammatory activation, harvested the cells and immediately proceeded to single cell RNA-seq library preparation, and construction of the guide-enrichment library through nested PCRs. Main- and guide enrichment libraries were subsequently pooled and sequenced on an Illumina Hi-seq machine to single end 50bp reads. To ensure adequate coverage and depth, we constructed two 10 000 cell 10X libraries from our edited cells and sequenced both to a depth of 50 000 reads per cell.

### CRISPR-Cas9 KO generation

KO cell lines of Kdm5c were generated in Cas9 positive ER-Hoxb8 cells. Guides targeting exon 2 of Kdm5c (Supplemental Table 3) were inserted into the pLKO5.sgRNA.EFS.GFP (Addgene, Plasmid #57822) backbone. Kdm5c guide plasmids were transfected into HEK293 cells plated in a T75 flask coated with Poly-L-Lysine. A virus assembly mix of 800 ng pMD2G, pMDL, pRSV each and 1600 ng plasmid was added to the HEK293 cells, together with 8 μl jetPRIME reagent and 400 μl jetPRIME buffer (jetPRIME Transfection Reagent, Polyplus). Medium was refreshed after 6 h and HEK293 cells were left to produce virus for 48 h. ER-Hoxb8 cells were transduced by adding 25% of virus medium and 8 ng/ml polybrene followed by spinoculation at 2000 rcf for 1 h RT. After 6 days, transduced cells were single-cell sorted on a SH800 Cell Sorter (Sony) into a 96 wells plate based on GFP intensity. Clones were expanded and KO status was confirmed by PCR and Sanger sequencing of the *Kdm5c* locus. Clones C1 and C24 had a frameshift resulting in a non-functional protein and were confirmed homozygous knockouts. Therefore, we selected C1 and C24 for downstream analyses.

### Western Blot

Nuclear protein fractions were extracted as described(24). Bolt Sample Reducing Agent (10x) (Novex) and Bolt LDS Sample Buffer (4x) (Novex) were added to the lysates and the mix was cooked at 95°C for 10 minutes. Bolt 4-12% Bis-Tris Plus (Invitrogen) gels were used in Bolt MOPS SDS running buffer (20x) (Invitrogen). The Spectra Multicolor High Range

Protein Ladder (Thermoscientific) was used as a reference. The gel was ran for 90 minutes. Using dry blotting, the bands were transferred to a Supported Nitrocellulose Membrane, 0.45 µm (Bio-Rad) for 30-45 minutes using Trans-Blot Turbo 5x Transfer Buffer (Bio-Rad). The membrane was blocked for an hour with 5% ELK + TBST. Primary antibodies (KDM5C: abcam ab194288, H3: Cell Signalling #4620S) were diluted 1:1000 in 5% ELK + TBST, and incubated overnight at 4°C.Next day, membranes were washed 10 minutes with 1% ELK + TBST, then exposed to 1:5000 secondary antibody in 5% ELK + TBST for 1 hour. After washing again for 3x 10 minutes in 1% ELK + TBST and 1x with TBS, the membrane was exposed 1:1 to the reagents of the SuperSignal West Pico PLUS Chemiluminescent Substrate for 10 minutes and imaged.

### RNA-seq

RNA was isolated and DNase treated using the RNeasy mini kit (Qiagen, #74106) following manufacturer’s instructions. mRNA-seq libraries were generated using the KAPA mRNA Hyperprep kit (Roche, #08098107702) and sequenced to a depth of 10-20 million paired-end 150 bp reads on an Illumina NovaSeq X Plus.

### Bioinformatic analyses

CROP-seq data were processed and analyzed using MUSIC, as described previously, to infer perturbation effects across transcriptional states(25).

Bulk RNA-seq data were mapped to mm39 using HISAT2. Count tables were constructed with HOMER’s anlayzeRepeats.pl and gene expression was analyzed using custom R scripts based on standard DESeq2 workflows for normalization, variance estimation, and differential expression analysis(26). Genes with an adjusted *p*-value < 0.05 were considered differentially expressed.

Single-cell RNA-seq data were analyzed using Seurat (v5) with standard preprocessing, normalization, integration, and clustering(27). Module scoring was performed using decoupleR(28): weighted mean enrichment (wmean) scores were computed per cell based on log□ fold-changes derived from the relevant bulk contrasts (e.g., *Kdm5c* KO vs WT). In this approach, each gene is assigned a directional weight proportional to its differential expression, and cell-wise enrichment is calculated as the weighted mean of normalized expression values across all genes in the module. Resulting scores were standardized and compared across cell clusters using linear mixed-effects models (random intercept = patient).

Transcription factor activity was inferred from the same single-cell dataset using DoRothEA(29) (confidence levels A/B) coupled to decoupleR’s univariate linear model (ULM) implementation to estimate per-cell regulon activity.

## Results

To identify histone modifying enzymes (HMEs) expressed in myeloid cells, we first analyzed gene expression profiles of all monocyte and macrophage samples available through the Blueprint consortium(30, 31). HMEs with RPKM > 1 were considered expressed (Supplemental Fig. 1A). We then expanded this list with HMEs differentially expressed between stable and unstable regions of human atherosclerotic plaques(32) (Supplemental Fig. 1B), resulting in 92 HMEs selected for inclusion in the CROP-seq library (Supplemental Table 1).

The pooled CRISPR library comprised three guides per HME, three proliferation-essential control genes (*Dhodh, Mvd, Tubb5*)(19), and twenty scrambled non-targeting controls (Supplemental Table 2). Lentiviral production yielded a pooled plasmid library with complete guide representation, as confirmed by sequencing (Figure 1A). Transduction of immortalized mouse bone-marrow-derived macrophages (BMDMs) followed by puromycin selection for 10 days preserved near-complete guide representation (Figure 1B). Comparison of guide abundances between the plasmid and transduced pools showed that non-targeting and HME guides remained close to their original levels, whereas the proliferation-essential controls were modestly but consistently depleted (∼2-fold), consistent with limited selective pressure in non-dividing macrophages (Figure 1C). Together, these data demonstrate even guide representation and efficient editing without severe bottleneck effects. Cells were then stimulated with 10 ng/ml LPS for 3 h, after which scRNA-seq and guide-enrichment libraries were generated.

**Figure 1.**
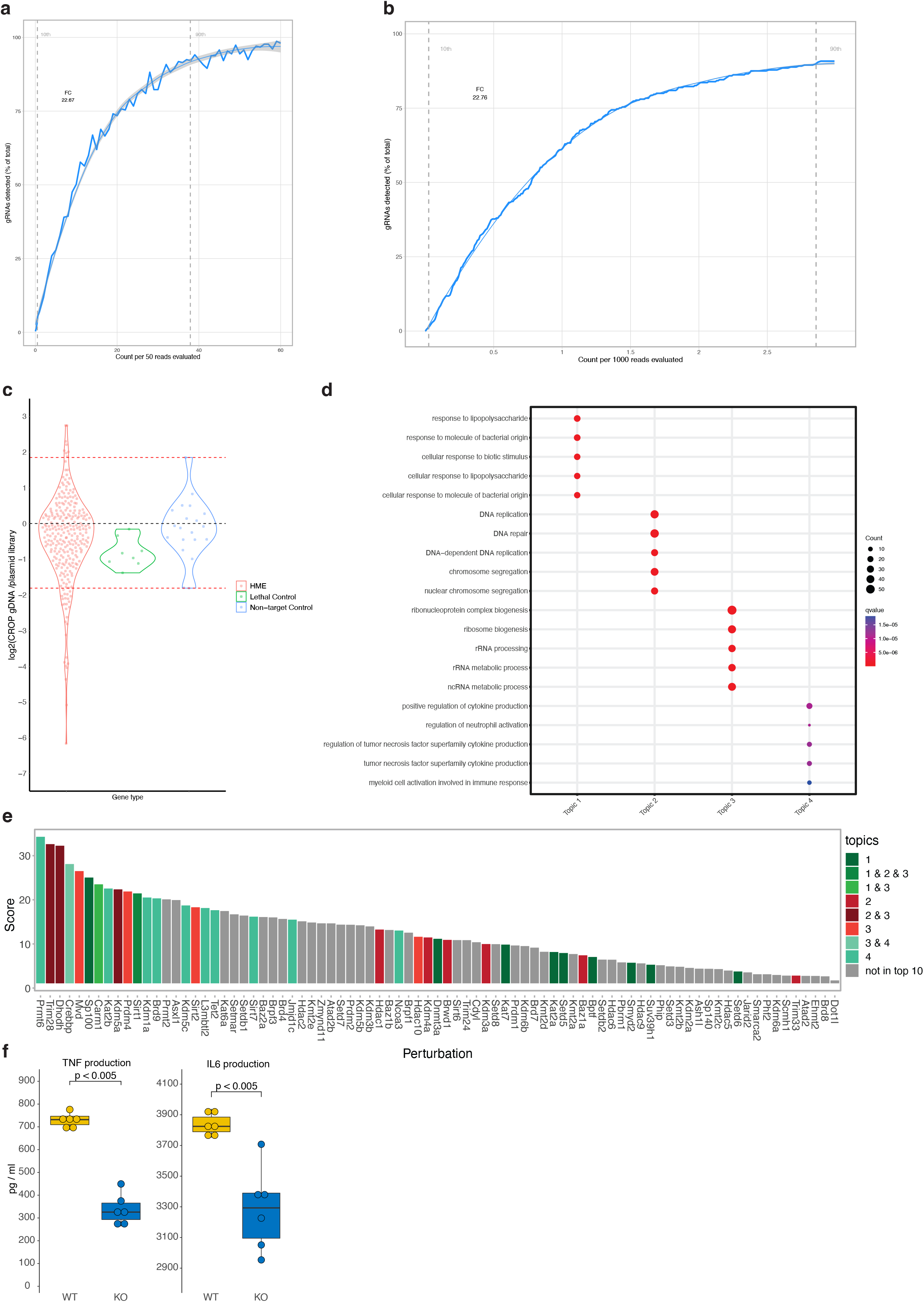
Inflammatory macrophage activation CROP-seq screen. (a) HME library guide RNA enrichment plots of the plasmid library and (b) transduced iBMDMs. (c) Violin plot of enrichment of guides in the plasmid library over the transduced iBMDMs. (d) Dot plot of enriched GO-terms in 4 topic clusters. (e) Bar plot of perturbation scores per HME. Colored by topic enrichment; the top 10 perturbations per topic are colored. (f) Boxplots of TNF (left) and IL6 (right) production in wild type (WT) or *Kat2b* deficient (KO) mouse peritoneal macrophages after 3 h of 10 ng/ml LPS stimulation, measured by ELISA.

After sequencing, single-cell transcriptomes were aligned to the *Mus musculus* reference genome (mm10) and linked to their corresponding perturbations by matching cellular barcodes with the paired guide-enrichment library. Guide assignment was successful for 92% of cells. Following quality control, we retained 15,652 high-quality cells that carried exactly one guide (Supplemental Figure 1C), expressed > 500 genes, > 1,000 unique molecular identifiers (UMIs), and contained < 20% mitochondrial transcripts (Supplemental Figure 1D).

Next, missing transcript counts were imputed using SAVER(33), and cells were filtered based on editing quality by comparing each cell’s gene-expression profile to the average profile of its assigned perturbation and scrambled controls. Only perturbations represented by at least 30 high-quality cells were retained for downstream analysis.

We then applied MUSIC(25) to infer latent transcriptional programs (“topics”) affected by each perturbation. The resulting topics captured two principal biological axes of variation: (1) inflammatory activation and (2) proliferation and cell-cycle control. These axes could be further resolved into four distinct topics: an LPS-response module (Topic 1), a DNA-replication module (Topic 2), a ribosome and RNA-metabolism module (Topic 3), and a cytokine-production / myeloid-activation module (Topic 4) (Figure 1D).

MUSIC then computed perturbation scores quantifying the extent to which each gene knockout modulated these topics, yielding a ranked list of HMEs according to their overall perturbation magnitude (Figure 1E).

Several of the top-ranked perturbations corresponded to established regulators of macrophage activation, confirming that the screen successfully captured perturbation-dependent transcriptional programs underlying inflammatory activation. Genes with strong effects on the LPS and cytokine topics (Topics 1 and 4) included chromatin-modifying enzymes and co-activators known to modulate the NF-κB pathway, such as *Prmt6(34), Carm1(35), Kat2b(36)*, and *Kdm1a(37)*. These hits cluster around transcriptional co-activator complexes that orchestrate inflammatory gene expression. For instance, *Prmt6* and *Carm1* are arginine methyltransferases that directly methylate the NF-κB p65 subunit, and both catalyze asymmetric dimethylation of H3R26 and H3R17 (H3R26me2a and H3R17me2a), although the precise contribution of these histone marks to inflammatory transcription remains unresolved. Another highly ranked hit, the histone acetyltransferase *Kat2b*, acetylates H3K9 and promotes pro-inflammatory transcription in macrophages, potentially acting in concert with PRMT6 and CARM1 by interaction with p300/CBP co-activator complexes(38). In contrast, perturbations loading on proliferation-associated topics (Topics 2 and 3) were enriched for chromatin regulators and metabolic enzymes linked to cell-growth and cell-cycle control, such as *Trim28(39), Kdm5a(40)*, and *Sirt1(41)*. Consistent with the MUSIC results, *Kat2b*-deficient mouse bone-marrow-derived macrophages displayed reduced inflammatory activation compared with wild-type controls (Figure 1F).

Another highly ranked hit was the histone demethylase Kdm5c, an X-linked member of the KDM5 family that removes tri- and di-methylation from histone H3 lysine 4 (H3K4me3/2)(42). Kdm5c can act both as a transcriptional activator, by erasing spurious H3K4me3 from enhancers to maintain proper chromatin architecture, and as a repressor, by demethylating promoter associated H3K4me3 at actively transcribed genes(43, 44). Through these dual functions, Kdm5c fine-tunes transcriptional amplitude rather than acting as a simple on/off switch and has been implicated in developmental(45, 46) and immune gene regulation(47, 48). In wild-type mouse thioglycolate-elicited peritoneal macrophages, *Kdm5c* expression increased after 3 h of LPS stimulation and was consistently higher in females, consistent with its X-linked dosage sensitivity (Figure 2A). The LPS-induced upregulation of Kdm5c supports its role as an active chromatin-level modulator of the inflammatory response.

**Figure 2.**
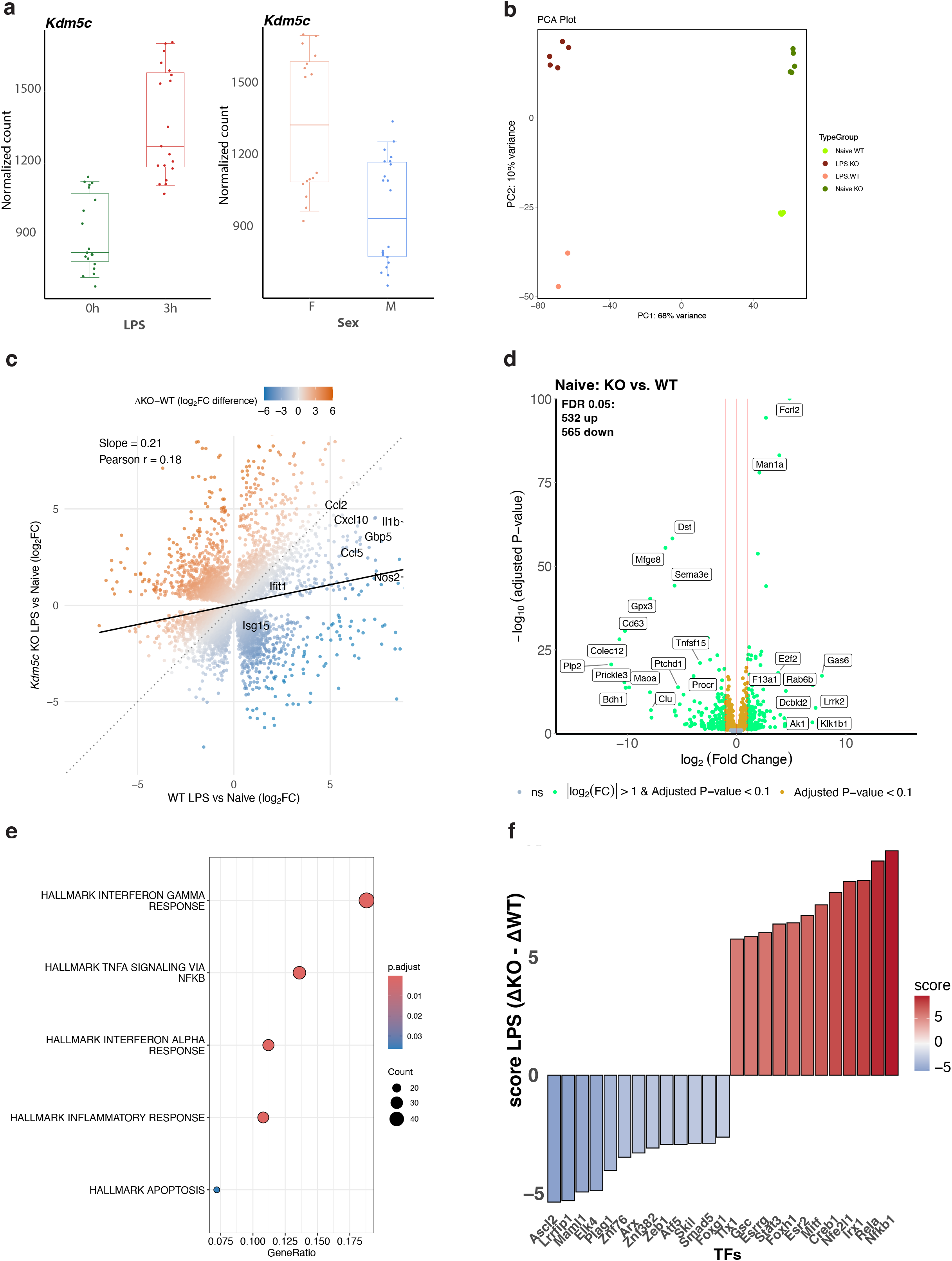
Kdm5c KO modulates the LPS response. (a) Boxplots of mouse peritoneal macrophages exposed to 10 ng /ml LPS for 3 h (left) or stratified by sex (right). (b) Bulk RNA-seq of *Kdm5c*-deficient (KO) and wild type (WT) ER-Hoxb8 mouse macrophages exposed to 3 h of 10 ng/ml LPS (LPS) or DMSO (Naïve). PCA Plot of experimental group distribution on principal component (PC) 1 and 2. (c) Scatter plot of LPS/Naïve log2(fold change) values in WT (x-axis) and KO (y-axis). Colors: ∂(KO-WT). Red = Up in KO. Blue = Down in KO. (d) Volcano plot of KO/WT log2(fold change) (x-axis) versus adjusted p-value (y-axis) in the naïve cells. (e) Dot plot of HALLMARK pathway overrepresentation analysis for genes downregulated in naïve KO cells versus naïve WT. (f) Bar plot of enrichment scores for transcription factor (TF) networks in KO versus WT cells after LPS treatment. Red = more induced in KO cells. Blue = more induced in WT cells.

To validate the effect of *Kdm5c* depletion on macrophage inflammatory activation, we generated a CRISPR–Cas9 induced knockout (KO) in ER-Hoxb8 mouse myeloid progenitor cells. Western blot analysis confirmed near-complete loss of the Kdm5c protein (Supplemental Fig. 2A). KO and wild-type (WT) ER-Hoxb8 cells were then differentiated into macrophages and stimulated with 10 ng/ml LPS for 3 h, after which the transcriptional response to *Kdm5c* loss was assessed by RNA-seq.

Samples neatly clustered on LPS stimulation along the first principal component and on *Kdm5c* status along the second (Fig. 2B) The LPS response was robustly triggered, with extensive transcriptional remodeling in both WT and *Kdm5c*-deficient macrophages (Supplemental Fig. 2BC). Direct comparison of the LPS responses revealed that *Kdm5c*-deficient macrophages deregulate the program overall (Pearson correlation = 0.18; Fig. 2C) and respond less strongly on average (slope = 0.21). Several canonical inflammatory genes, including *Ccl2, Cxcl10*, and *Nos2*, showed reduced induction in the KO.

In unstimulated macrophages, *Kdm5c* knockout altered the expression of 1,097 genes at FDR < 0.05 (Fig. 2D), characterized by broad downregulation of inflammatory and immune-activation related HALLMARK pathways, e.g., ‘INTERFERON GAMMA RESPONSE’, ‘TNFA SIGNALLING VIA NFKB’, ‘INFLAMMATROY RESPONSE’ (Fig. 2E).

When comparing the magnitude of the LPS response between genotypes using an interaction model (ΔKO − ΔWT), inflammatory programs emerged among the most strongly enriched in the knockout (Supplemental Figure 2D), as exemplified by impact on gene set enrichment analysis (Supplemental Figure 2E) and transcription factor (TF) network expression (Figure 2F). E.g., Rela and Nfkb1TF networks and their corresponding pathways like ‘TNFA SIGNALLING VIA NFKB’ are enriched in the KO setting. This indicates that while the induction of individual inflammatory genes was somewhat blunted, TF network and pathway level analyses point to a broader shift toward heightened inflammatory responsiveness in *Kdm5c*-deficient macrophages. Because inflammatory pathways were suppressed at baseline, we next tested whether the higher inducibility in *Kdm5c*-deficient macrophages simply reflected a “catch-up” from this repressed state or represented a true overshoot. Direct comparison of LPS-stimulated KO and WT macrophages showed that inflammatory pathways such as ‘INFLAMMATROY RESPONSE’ and ‘TNFA SIGNALLING VIA NFKB’ remained enriched in the KO setting (Supplemental Figures 2EF).

Together, these results indicate that Kdm5c restrains macrophage inflammatory gene expression at baseline but enhances the amplitude of the acute LPS response, consistent with a model in which Kdm5c modulates chromatin accessibility to balance tonic and inducible transcriptional activity through its dual promoter-silencing and enhancer-activating functions.

Lastly, to explore the relevance of these findings in human disease, we analyzed single-cell RNA-seq data from human atherosclerotic plaque macrophages(16) to explore the role of KDM5C in a human disease setting. Here, we found that *KDM5C* is mostly expressed in inflammatory macrophages (Fig. 3A) and, consistent with its X-linked dosage sensitivity, was more strongly expressed in females (Supplemental Fig. 3A). Weighted module scoring of the *Kdm5c*-deficient macrophage signature from the KO/WT bulk RNA-seq also revealed an enrichment of the KO signature in inflammatory macrophages (Fig. 3B).

**Figure 3.**
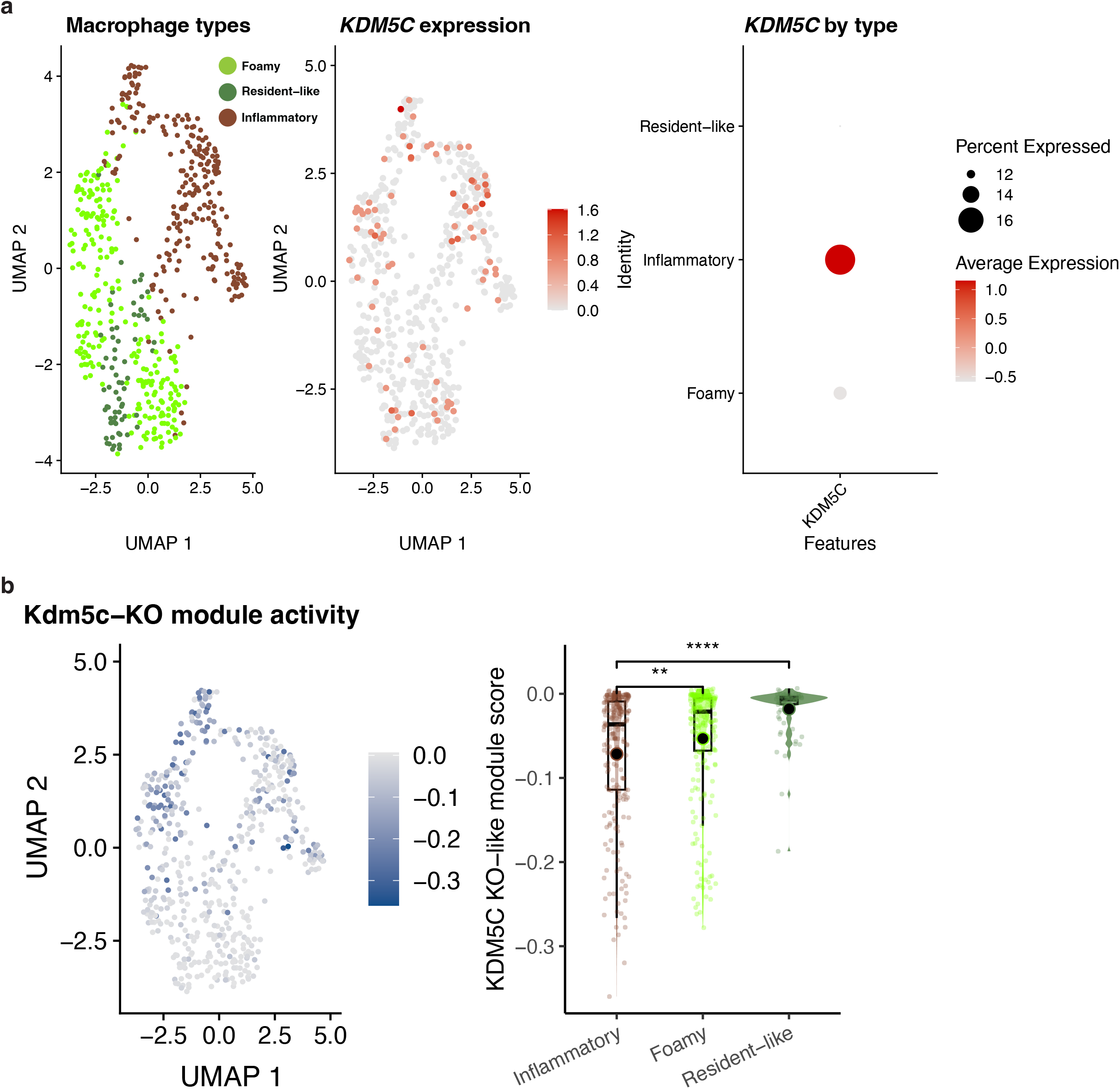
KDM5C activity in human atherosclerotic plaque macrophages. (a) UMAPs of macrophage population identities (left) and *KDM5C* expression (middle). Dot plot of *KDM5C* expression z-score (color) and abundance (size) per population (right). (b) UMAP (left) and bar plot (right) of weighted *Kdm5c* module-score in human plaque macrophages. Negative enrichment values denote a KO-like phenotype.

Together, these findings suggest that the inflammatory macrophage phenotype may, at least in part, reflect transcriptional programs normally restrained by KDM5C and downstream TLR4/NF-κB–STAT3 signaling.

## Discussion

This study demonstrates the power of CROP-seq to dissect the epigenetic regulation of macrophage activation. By coupling pooled CRISPR screening to single-cell transcriptomics, we systematically evaluated the contribution of 92 histone modifying enzymes (HMEs) to inflammatory gene regulation in macrophages. The resulting dataset captures a broad regulatory landscape, confirming known modulators and revealing less characterized enzymes that influence inflammatory tone.

Among the top hits were *Prmt6, Carm1, Kat2b*, and *Kdm5c*. The identification of *Prmt6* and *Carm1* is particularly noteworthy, as both are protein arginine methyltransferases that catalyze asymmetric dimethylation of arginine residues on histones and transcriptional regulators—modifications less extensively characterized in macrophages than the canonical lysine marks. Prmt6 and Carm1 methylate both histone H3 (e.g., H3R2me2a, H3R17me2a, H3R26me2a) and non-histone substrates such as the NF-κB p65 subunit, thereby influencing enhancer accessibility and co-activator recruitment(34, 35). Notably, both enzymes have been reported to associate with the CBP/p300 co-activator complex(38), placing them in the same regulatory context as *Kat2b*, which acetylates H3K9 to promote transcriptional activation(36). Thus, multiple top hits from our screen converge mechanistically on the same CBP/p300-associated axis that coordinates enhancer acetylation and arginine methylation to control inflammatory gene expression. Future studies should examine these interactions directly, for example by combining individual knockouts with ChIP-seq or CUT&Tag profiling for H3Rme2a and H3K9ac to define overlapping enhancer signatures and potential co-occupancy within shared co-activator complexes.

Our detailed validation of *Kdm5c* further highlights the complexity of epigenetic control in macrophage activation. Loss of *Kdm5c* reduced basal inflammatory gene expression but enhanced inducibility upon LPS stimulation, suggesting that Kdm5c fine-tunes transcriptional amplitude rather than acting as a simple repressor. This dual behavior likely reflects its known ability to demethylate H3K4me3 at both promoters and enhancers, balancing chromatin accessibility and signal responsiveness. Follow-up studies should assess genome-wide redistribution of H3K4 methylation and accompanying transcription factor occupancy in *Kdm5c*-deficient macrophages, ideally integrating ChIP-seq for H3K4me1/3 and H3K27ac with ATAC-seq to analyze enhancer and promoter accessibility changes and transcription factor motif enrichments. Functional validation in human monocyte-derived macrophages would further clarify whether this regulatory mechanism is conserved across species and sexes.

Finally, our single-cell analysis of human atherosclerotic plaques shows that *KDM5C* expression and its downstream transcription factor networks are enriched in inflammatory macrophages. While correlative, this finding raises the possibility that KDM5C contributes to the regulation of macrophage-driven inflammation and plaque instability. Future studies could examine whether modulating KDM5C activity—or its downstream effectors—offers a means to temper chronic inflammation in atherosclerosis without broadly suppressing macrophage function.

## Limitations

Our study has several limitations. The CROP-seq screen was performed in immortalized mouse macrophages under acute LPS stimulation, which may not fully recapitulate the epigenetic landscape of primary or tissue-resident macrophages. We also inferred chromatin-level mechanisms from transcriptional data without direct histone modification profiling, so future experiments integrating ChIP-seq, CUT&Tag, or ATAC-seq will be necessary to define the precise genomic targets of KDM5C and other HMEs. Finally, while our cross-species comparison with human plaque macrophages supports translational relevance, functional studies in primary human systems and in vivo models of atherosclerosis will be needed to establish causal relationships between KDM5C activity, macrophage phenotype, and plaque stability.

## Conclusion

In summary, this work establishes a scalable framework to explore the epigenetic control of macrophage activation and identifies *Kdm5c* as a key regulator of the balance between tonic and inducible inflammatory gene expression, with potential implications for inflammatory disease and vascular pathology.

## Supporting information

Supplemental Table 1 (HMEs)

Supplemental Table 2 (guide library)

Supplemental Table 3 (Kdm5c guides)

## Acknowledgements

Menno de Winther is supported by the Netherlands Heart Foundation (AtheroNeth, 01-001-2024-0601, a collaboration supported by the Dutch CardioVascular Alliance (DCVA)); Leducq Foundation (LEAN Transatlantic Network Grant, 16CVD01); Amsterdam Cardiovascular Sciences; ZonMW (Open competition 09120011910025) and the European Union (Horizon Europe research and innovation program under Marie Sklodowska-Curie Actions Doctoral Network program 2022 under Grant Agreement No. 101119370; MIRACLE). Koen Prange is supported by a Veni grant from the Dutch Research Council (NWO) (VI.VENI.212.196).

We thank Dr. P. Quax (Leiden University Medical Center) for the generous gift of Kat2b deficient bone marrow.

## Figure Legends

**Supplemental Figure 1:**
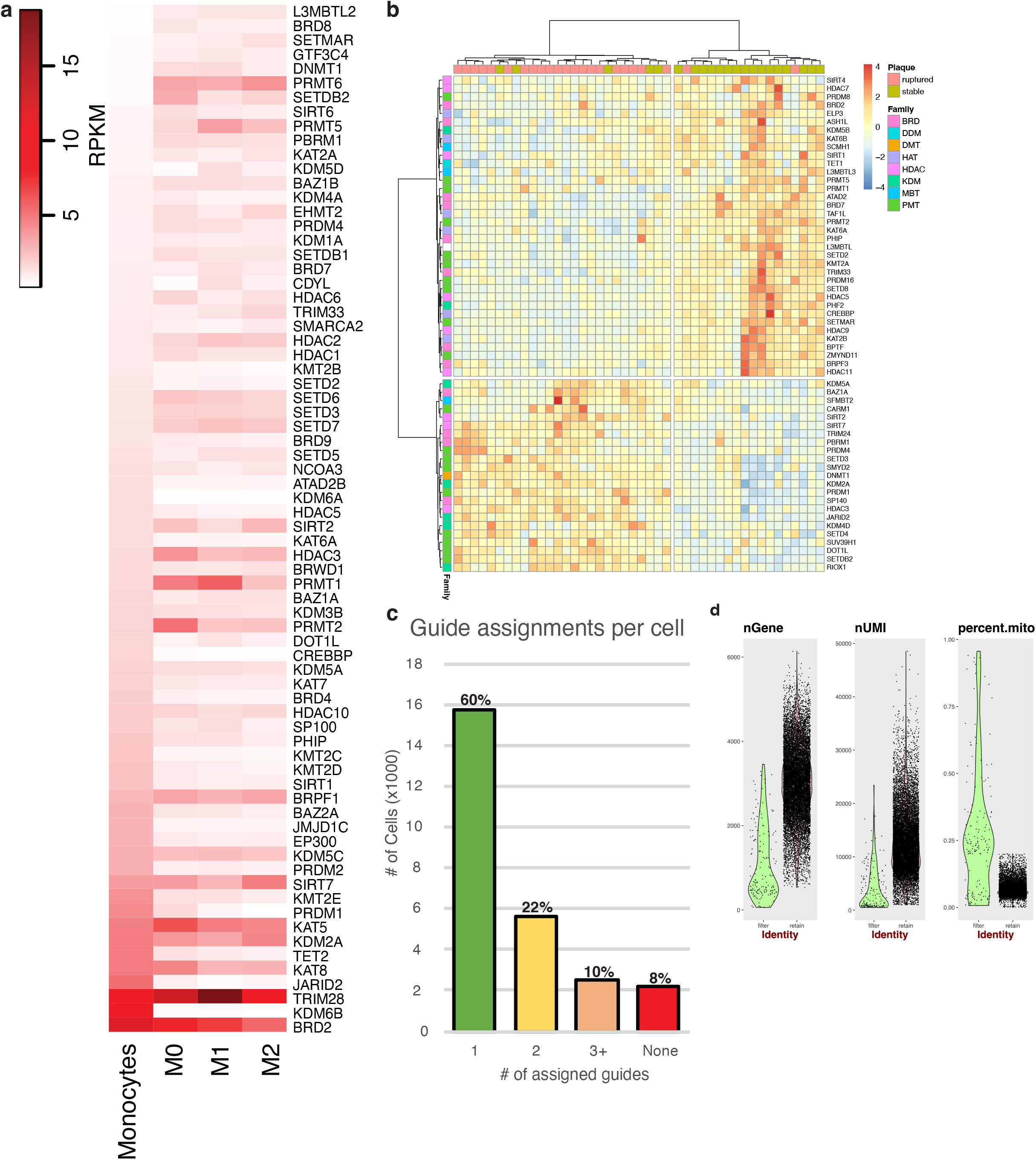
Inflammatory macrophage activation CROP-seq screen. (a) Heatmap of monocyte and macrophage expression in the Blueprint dataset. RPKM = reads per kilobase per million mapped reads. (b) Heatmap of z-scores of HME expression in stable versus ruptured regions of human atherosclerotic plaques as measured by micro-array. (c) Bar plot of CROP-seq guide assignment distribution in the scRNA-seq libraries. (d) Quality control violin plots of the scRNA-seq libraries showing the number of genes per cell (left), number of UMIs per cell (middle), and the percentage of mitochondrial reads per cell (right).

**Supplemental Figure 2:**
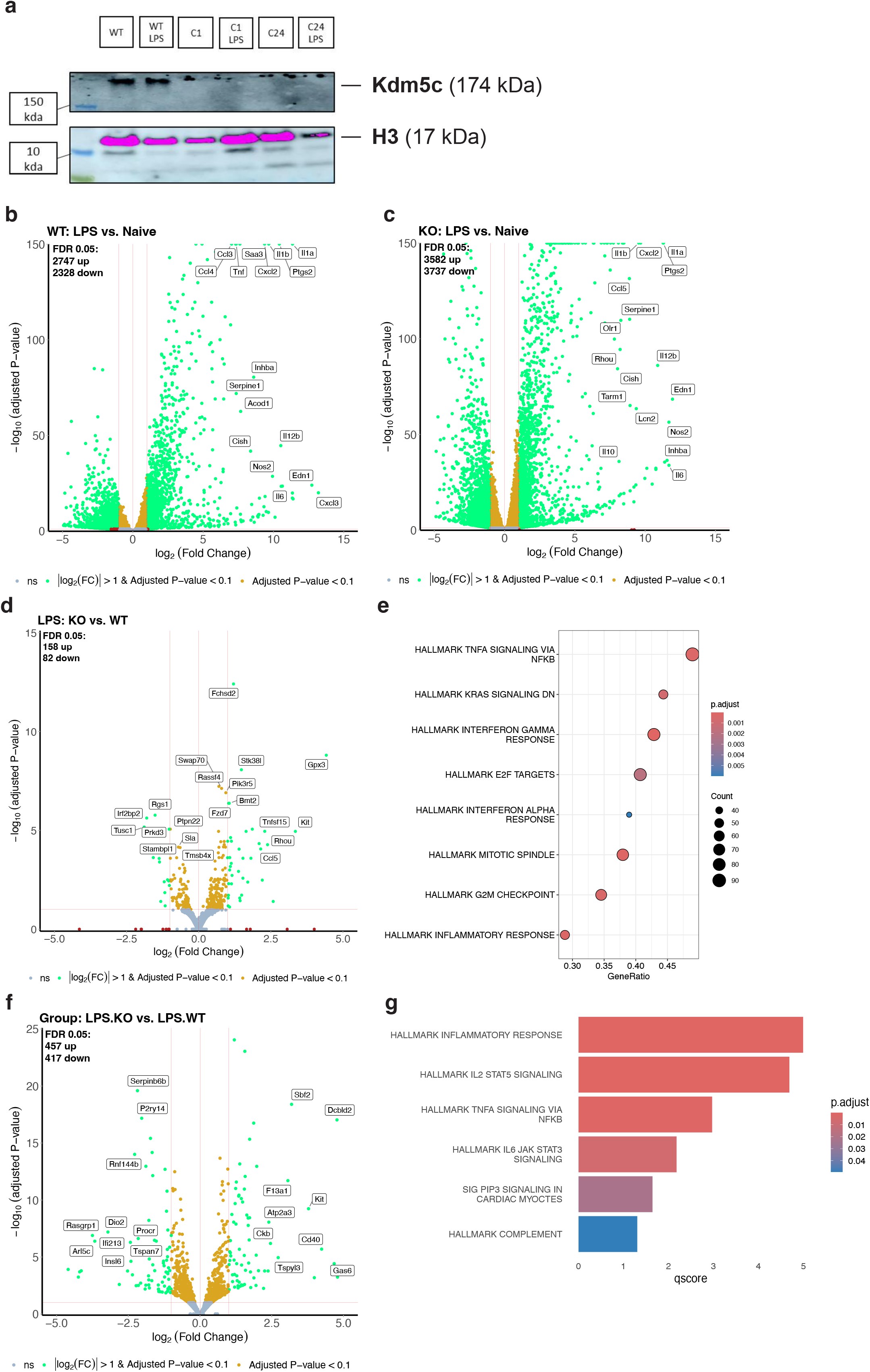
Kdm5c KO modulates the LPS response. (a) Gel image of western blot using Kdm5c and H3 antibodies in WT and *Kdm5c*-deficient ER-Hoxb8 macrophages. WT = wild type, C1 = KO Clone 1, C24 is KO Clone 24. kDa = kilodalton. (b) Volcano plot of LPS/Naïve log2(fold change) (x-axis) versus adjusted p-value (y-axis) in the WT cells. (c) Volcano plot of LPS/Naïve log2(fold change) (x-axis) versus adjusted p-value (y-axis) in the KO cells. (d) Volcano plot of the interaction term (∂KO - ∂WT) of KO/WT log2(fold change) (x-axis) versus adjusted p-value (y-axis) in the LPS cells. (e) Dot plot of HALLMARK gene set enrichment analysis for genes more induced in KO cells versus WT in an LPS setting (∂KO - ∂WT). (f) Volcano plot of per group KO.LPS/WT.LPS log2(fold change) (x-axis) versus adjusted p-value (y-axis).

**Supplemental Figure 3:**
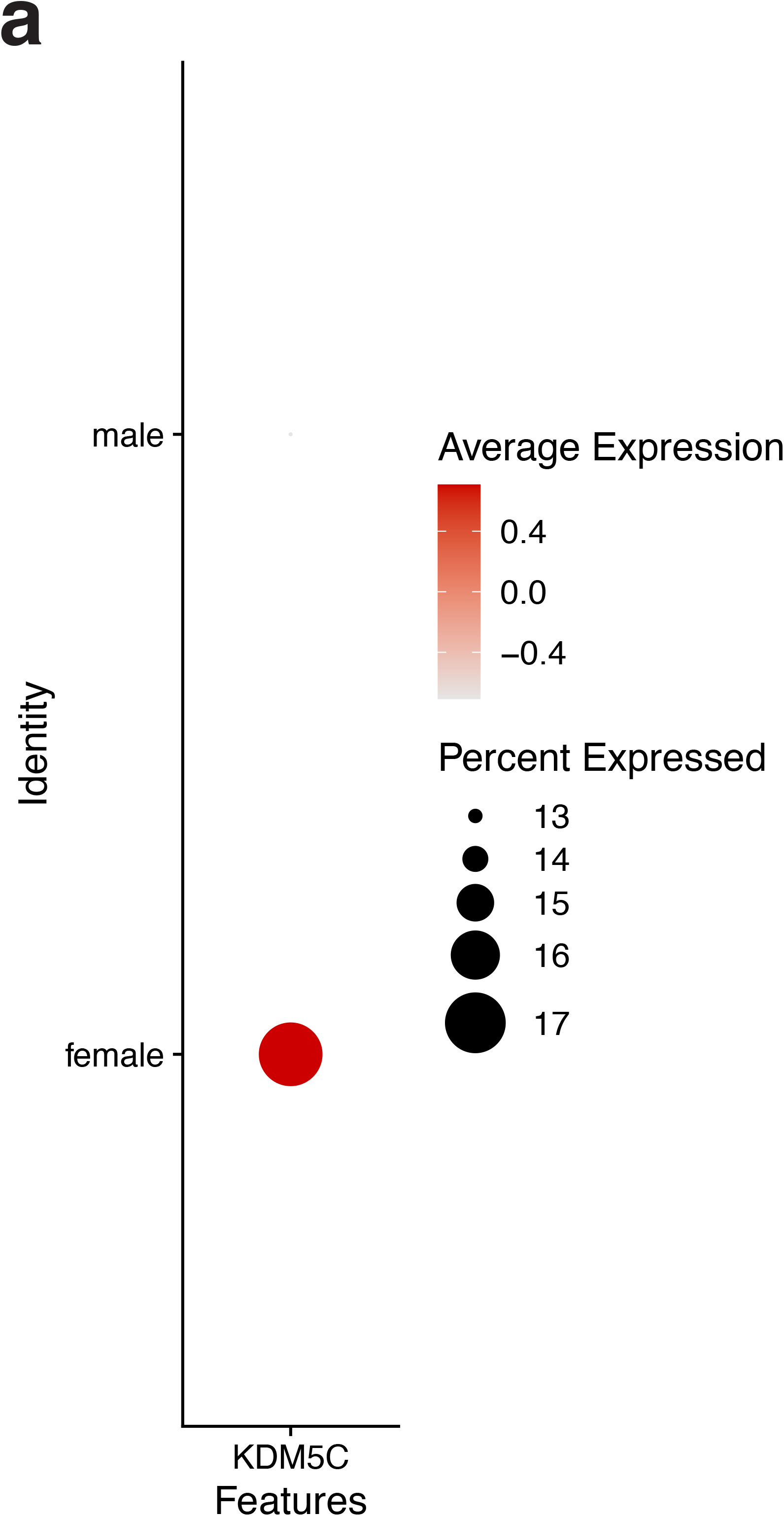
KDM5C activity in human atherosclerotic plaque macrophages. (a) Dot plot of *KDM5C* expression z-score (color) and abundance (size) in human atherosclerotic plaque macrophages stratified by sex.

